# Multi-task benchmarking of spatially resolved gene expression simulation models

**DOI:** 10.1101/2024.05.29.596418

**Authors:** Xiaoqi Liang, Yue Cao, Jean Yee Hwa Yang

## Abstract

Computational methods for spatially resolved transcriptomics (SRT) are frequently developed and assessed through data simulation. The effectiveness of these evaluations relies on the simulation methods’ ability to accurately reflect experimental data. However, a systematic evaluation framework for spatial simulators is lacking. Here, we present SpatialSimBench, a comprehensive evaluation framework that assesses 13 simulation methods using 10 distinct STR datasets. We introduce simAdaptor, a tool that extends single-cell simulators to incorporate spatial variables, thus enabling them to simulate spatial data. SimAdaptor enables SpatialSimBench to be “back-wards” compatible. That is, it facilitates direct comparison between spatially aware simulators and existing non-spatial single-cell simulators through the adaption. Through SpatialSimBench, we demonstrate the feasibility of leveraging existing single-cell simulators for SRT data and highlight performance differences among methods. Additionally, we evaluate the simulation methods based on a total of 35 metrics across data property estimation, various downstream analysis and scalability. In total, we generated 4550 results from 13 simulation methods, 10 spatial datasets and 35 metrics. Our findings reveal that model estimation can be impacted by distribution assumptions and dataset characteristics. In summary, our evaluation and the evaluation framework will provide guidelines for selecting appropriate methods for specific scenarios and informing future method development.

## Introduction

Spatial transcriptomics (ST) technology represents a significant advancement in the field of molecular biology, offering the ability to map gene expression data within the spatial context of tissue samples[1]. In many situations, the ground truth such as differential expression or differential abundance gene is experimentally unattainable. Simulation, in contrast, provides access to a controlled environment with ground truth, thereby enabling systematic evaluation of algorithms. Spatial simulations play an essential role in validating the efficacy of computational tools such as CARD[2], stLearn[3], SPIAT[4], and BayesSpace[5]. These tools address a range of challenges including cell type deconvolution, cell-cell interaction analysis, tissue microenvironment characterization, and sub-spot resolution enrichment.

Hence, computational simulation methods that generate spatial datasets stand as a crucial strategy for assessing the performance of spatial analytical tools.

In a recent study, simBench[6] has recognized the importance of simulated datasets for methodology development and provides a platform for assessing how well the various simulation tools reflect real-world data. With the increased demand of spatial analytical tools, there is an emerging but limited number of spatially aware simulators developed to assist with methods development. We can categorize spatially aware simulators based on the type of input: spot-level data (as in Visium ST) and scRNA-seq data. Spot-level data simulators, such as scDesign3 and SRTsim, generate spot-level count data while preserving the spatial layout observed in real data. scDesign3 focuses on reference-based simulations, whereas SRTsim can handle both reference-based and reference-free scenarios.

Another category utilizes scRNA-seq data as input. Simulators like spider[7], stLearn[3], and SpatialcoGCN-sim[8] fall into this category, generating spot-level count data and spatial location. While some scRNA-seq based simulators, such as spaSim[4], solely focus on cell location simulation, others like scMultiSim[9] and stLearn[3] can simulate both cell counts and spatial cell-cell interaction relationships. It is important to note that some simulators used in publication such as CARD[2], are not published independently but rather used as part of the evaluation process for published methods.

Considering the large number of single-cell simulators currently available and the relatively few spatially aware simulators that accept spot-level data as input, it is essential not to overlook existing tools or to start from scratch when developing spatial simulators. One might investigate the opportunities to adapt the capabilities of current single-cell simulators to simulate for spatially resolved data. These single-cell simulators, as highlighted in the simBench[6] study, includes SPARsim[10], ZINB-WaVE[11], Splatter[12], and SymSim[13].

To this end, our study, SpatialSimBench, is the first single-cell benchmarking study to examine simulation approaches with specific attention to spatial expression. In particular, we address the opportunity of leveraging existing single cell simulators and develop simAdaptor, a tool that enables the extension of single-cell simulators to simulate spatial data by incorporating spatial variables. We have devised different simulation strategies to make the benchmarking “back-wards” compatible. That is, we examine in one frame between (i) spatially aware simulation models and (ii) existing “non-spatial” simulation methods that are adapted. Our benchmarking design will leverage and extend the framework developed in our previous work, simBench[6]. It will (i) examine both two input types of spatially aware simulators methods; (ii) introduce spatially specific metrics to examine data properties (simBench[6]) and (iii) examine impact on multiple downstream analysis tasks that is typically done in spatially analysis. Finally, we compile the findings into recommendations for users and emphasize potential areas for future research.

## Results

### Leverage existing scRNA-seq simulation for spatial resolved data to capture spatial patterns

To examine whether we can leverage the large collection of existing single-cell simulations for spatial simulation, we develop a simAdaptor method that incorporates spatial variables into single-cell simulators. The strategy here is to first employ spatial clustering to identify groups of regions with similar gene expression profiles. This approach relies on the assumption that distinct spatial clusters will harbor transcriptional features within their respective regions. In the initial step, clusters are manually created with the assumption that each cluster represents a distinct spatial region. Following this, each cluster is then utilized as input for spatial or single-cell simulation models to simulate individual spatial regions. We term this approach as simAdaptor which uses regional information as the foundation.

To illustrate the efficacy of the simAdaptor, we generated spatial simulation data using an adult mouse olfactory bulb spatial gene expression data as the reference (Fig. 2). Figure 2a demonstrates the initial spatial clustering of the data into four distinct groups (Fig. 2a). Differential gene expression analysis is conducted within individual spatial clustering groups to highlight distinct gene expression patterns associated with each group. Following this, five single-cell simulators were employed to assess their ability to capture data distribution of the spatial data. From Figure 2b,c, the simulation data from scDesign2, SPARsim, and ZINB-WaVE show similar distribution compared with real data particularly in gene-level and spot-level. In spatial-level evaluation, scDesign2, SPARsim, Splatter outperformed others (Fig. 2d). In evaluating the performance on the adapted simulator in identifying regional structure, we found the adapted methods based on SPARsim, Splatter, and SymSim show consistent spatial clustering patterns with real data (Fig. 3a). Similarly, SPARsim, Splatter and ZINB-WaVE effectively captured the majority of cell type proportions observed in the real data (Fig. 3b). In spatial autocorrelation, scDesign2 and SPARsim outperformed (Fig. 3c). SPARsim and Splatter performed well in selected spatially variable genes (Fig. 3d). Our analysis of spatial patterns revealed that specific single-cell simulators, including SPARsim, Splatter, ZINB-WaVE and SymSim, generated spatial patterns consistent with those observed in the real spatial data. Additionally, spot-spot relationships, such as cell-type interactions and Moran’s I, exhibited similar distributions to the real data. These findings demonstrate the effectiveness of our approach in adapting single-cell simulators for generating data with spatial features (Fig. 2d).

**Figure 1:**
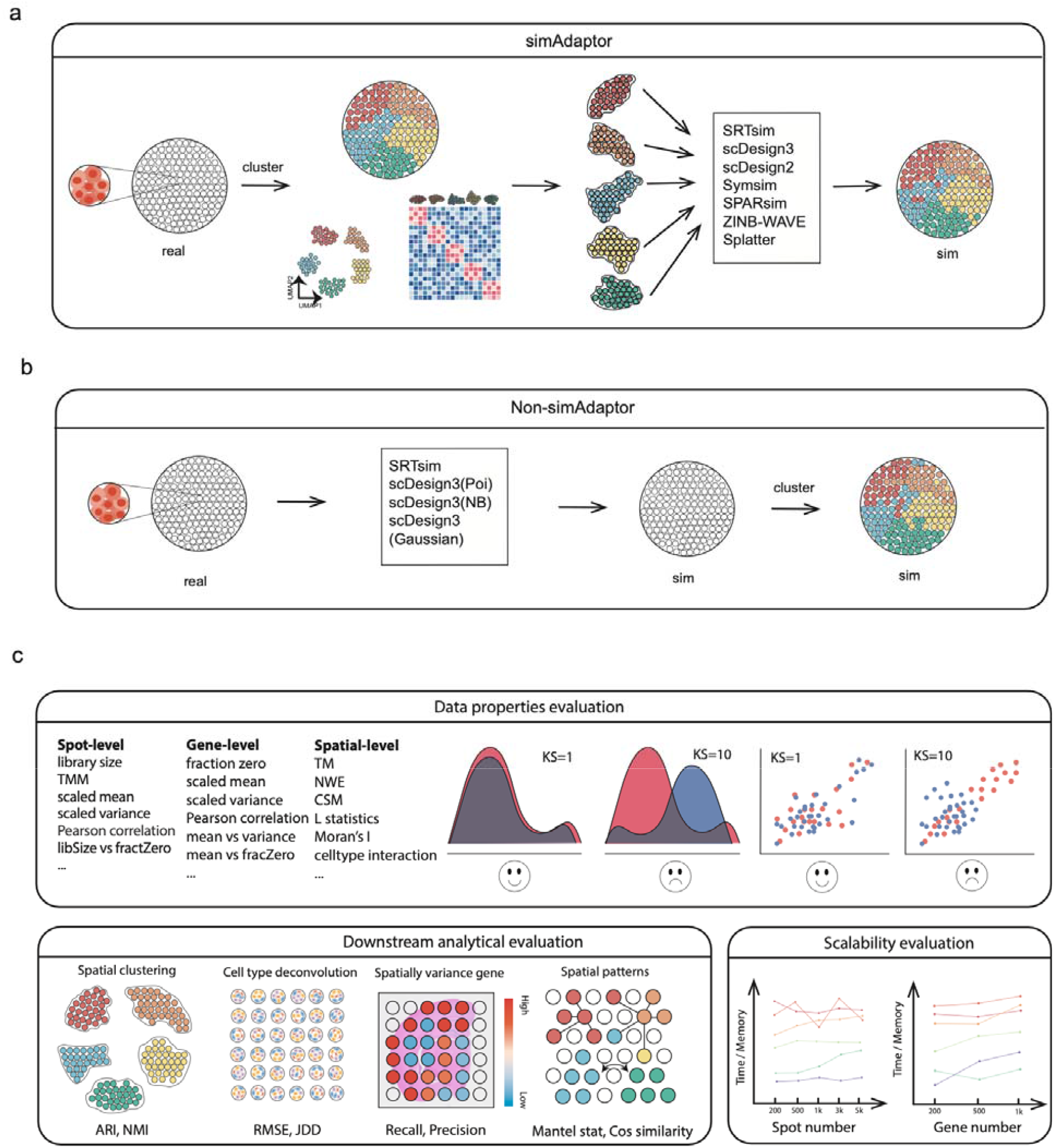
Overview of the benchmarking process and key aspects of evaluation. **a**, Schematic of simAdaptor approach. It applies spatial simulated models directly without segmentation considerations. **b**, Schematic of non-simAdaptor approach. Initially, spatial clustering of data identifies regions sharing similar expression profiles. Following this, spatial transcriptomics data is categorized, allowing the application of established simulation methods to each identified category. **c**, Multi-tasks of evaluation were examined in this study, including data properties, spatial downstream analytical task and scalability.

**Figure 2:**
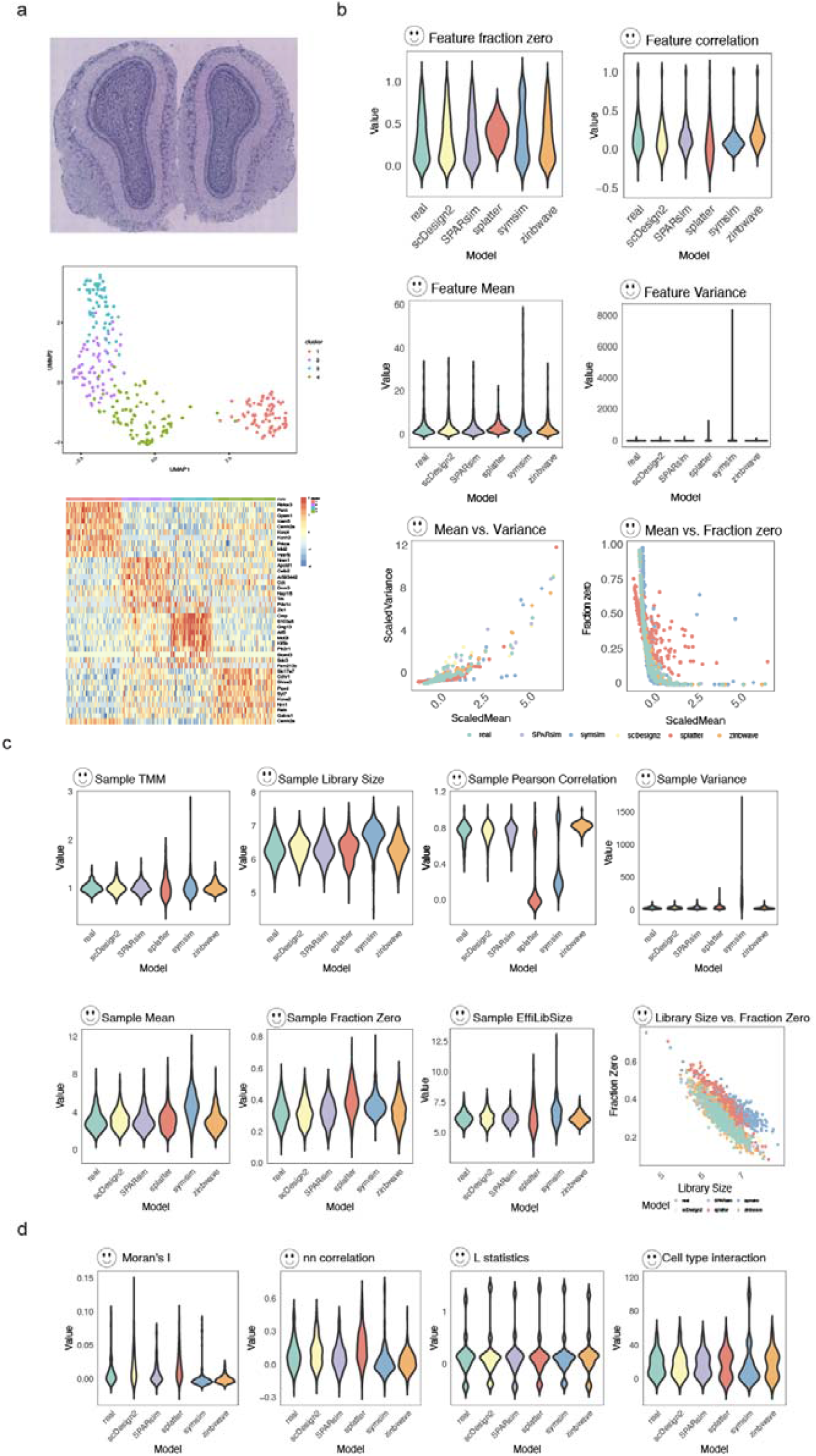
Overview of the simAdaptor approach with data properties evaluation result. **a**, Performance spatial clustering and differential expression analysis on real data **b**, Visualize the real and simulation in boxplot across gene-level metrics. **c**, Visualize the real and simulation in boxplot across spot-level metrics. **d**, Visualize the real and simulation in boxplot across spatial-level metrics.

**Figure 3:**
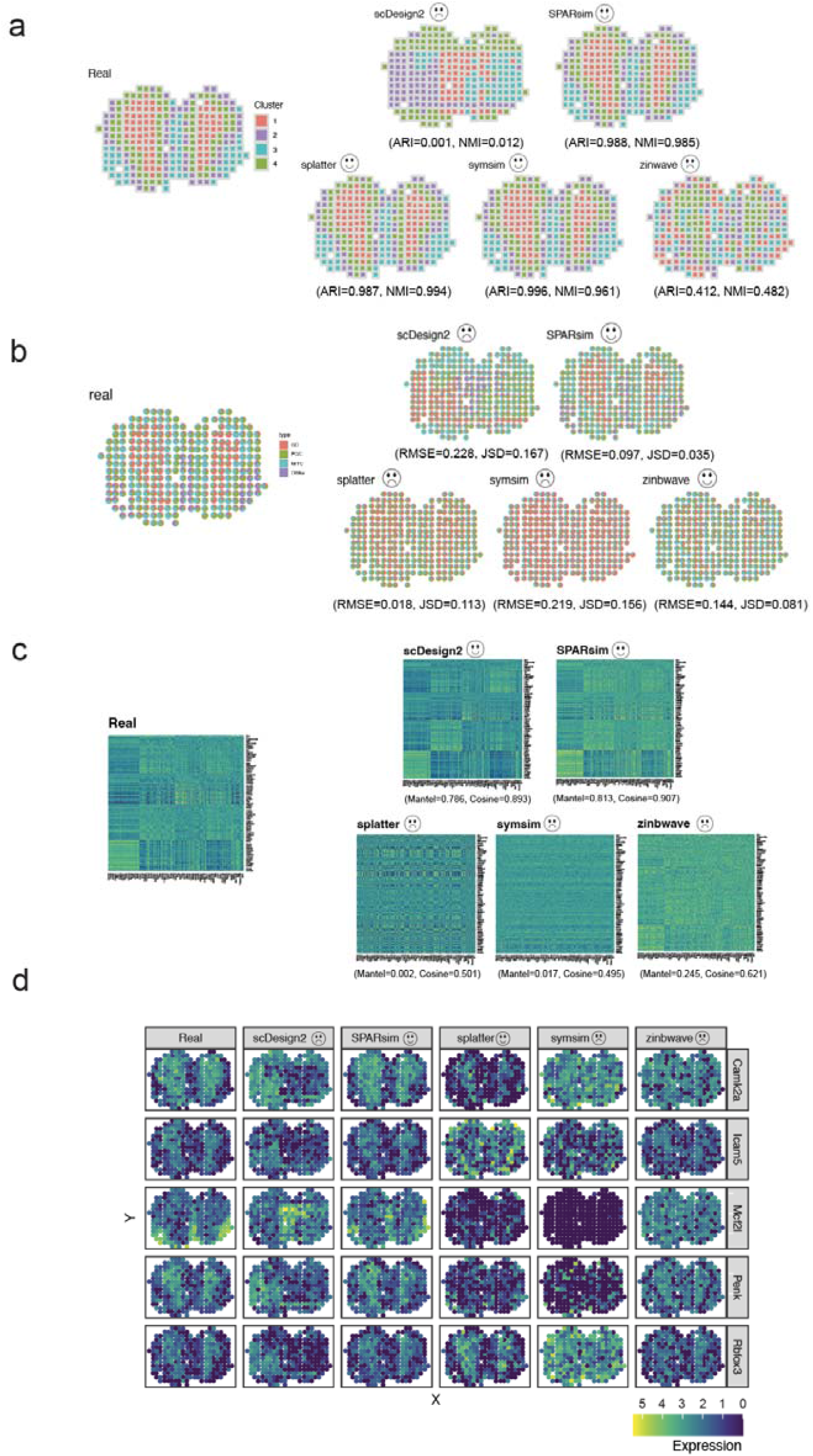
Overview of the simAdaptor approach with spatial downstream analysis result. **a**, Spatial clustering visualization comparison. **b**, cell type deconvolution visualization comparison. **c**, Spatial cross correlation visualization comparison. **d**, Selected spatial variance gene visualization comparison.

Next, we assess the applicability of the simAdaptor approach on four spatially aware simulators and whether this approach will improve their performance (Supplementary Fig. 1). Visualizations across the spot-level, gene-level, and spatial-level metrics revealed a high degree of similarity between the simAdpator and non-simAdpator, suggesting compatibility of simAdpator approach with both spatial and single-cell simulators. Notably, our findings indicate that the simAdpator approach led to improvements in specific performance metrics for spatial simulators, particularly in spot-wise scaled mean and variance, as shown in Figure 4b.

**Figure 4:**
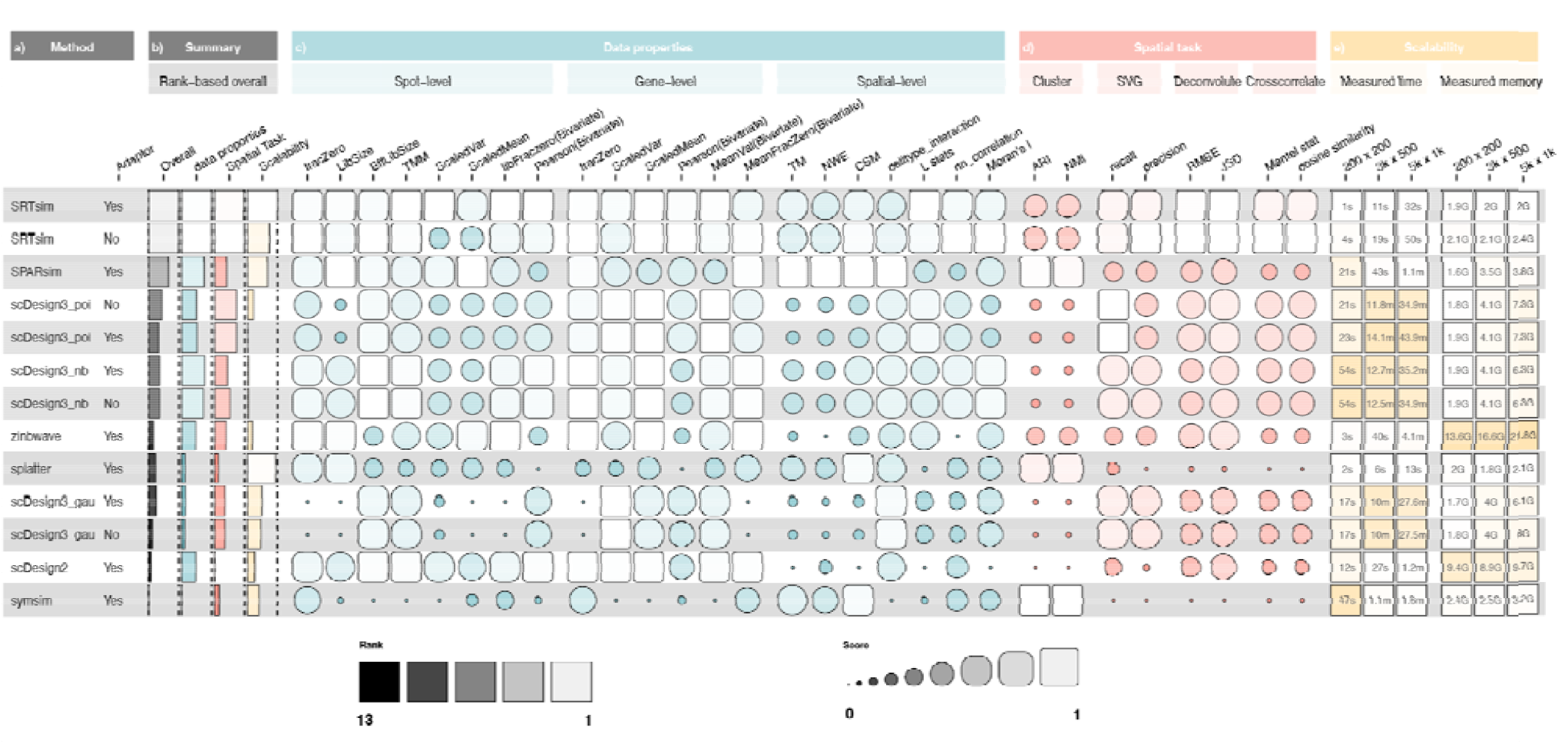
Details result of three main evaluation metrics: data properties, spatial task and scalability. The colour represents different areas of evaluation and the higher score shows the best possible rank of 1. **a**, The name of the method across Non-simAdaptor and simAdaptor approaches. **b**, summary of all the overall performance **c**, Score of methods within data properties, ranking by KDE value. **d**, Score of methods within spatial tasks, ranking by KDE value. **E**, Scalability results for varying numbers of spots and features (number of spots x number of features). K, thousands.

### SpatialSimBench is a comprehensive benchmark of spatially resolved simulation methods using diverse datasets and comparison measures

Our SpatialSimBench framework evaluates recently published spatially aware simulation methods together with single-cell simulation methods adapted with simAdaptor (Fig. 1a, Table 1). This SpatialSimBench includes a total of 35 metrics that comprehensively examine data property estimation (spot-level, gene-level and spatial-level), diverse range of spatial tasks (clustering, spatially variable gene identification, cell type deconvolution and spatial cross-correlation) as well as scalability. In our previous study, we introduced simBench, a benchmark study for simulation methods for scRNA-seq data. We leverage the various categories of gene and cell level properties and scalability developed in simBench and introduce two additional categories of spatial specific metrics. The first category refers to the simulator’s ability to capture data properties of spot level and spatial level and the second focuses on the simulator’s performance capacity in various spatial downstream tasks. Similar to simBench, to ensure robustness and generalisability of the study results and account for variability across datasets, we collect 10 public spatial transcriptomics experimental datasets, encompassing a variety of sequencing protocols, tissue types, and health conditions from human and mouse sources. Spatial simulation data were generated by using these real experimental datasets as reference. The simulation was then assessed against the real data using the three aspects of metrics (Fig.1c). Through this systematic comparison, we generated a total of 4550 results derived from 10 spatial datasets, 13 simulation methods and 35 metrics.

To assess the similarity between a real dataset and simulated dataset in data properties, we use both cell-level (spot-level in spatial data) and gene-level metrics, based on our previous benchmarking study on single-cell simulators[6]. This involved defining metrics encompassing various aspects, including spots density, along with higher-order interactions, such as mean-variance relationships. To capture the spatial dimension, we evaluate how well each simulator captures spot-spot (or cell-cell) relationships in the spatial setting, which is called spatial-level in the data properties section. This is achieved by analyzing both real and simulated data with transition matrices, neighborhood enrichment matrices, and centralized score matrices to quantify spatial relationships between spots (see Methods). Also, we added metrics (cell type interaction, L statistics, nn correlation and Moran’s I) from our previous study called scFeatures[14], multi-view representations of single-cell and spatial data at the spot level. To evaluate the similarity between simulated and real data for each metric, we utilized two methods: 1) density plots for visual inspection and 2) a kernel density-based global two-sample comparison KED test statistic[15] for quantitative assessment. To assess the efficiency of our simulation models in generating large-scale datasets, we investigated computational scalability. This involved measuring the simulation models’ running time and memory usage while simulating datasets with varying sizes of spots and genes.

We also propose spatial aware metrics to evaluate a simulation model’s ability in spatial patterns, cell type composition, spatially variable genes identification and spatial cross-correlation of real data. To examine how the simulator is able to capture some of the standard downstream analysis, we focus on downstream tasks. These tasks include: 1) maintaining spatial clustering, measured by Adjusted Rand Index (ARI) and Normalized Mutual Information (NMI); 2) performing cell-type deconvolution, where simulated data is compared to real data using the same deconvolution algorithm and evaluated using Root-mean-square deviation (RMSE) and Jensen-SHannon divergence (JSD); 3) the ability to accurately identify spatially variable genes (SVG), where simulated data is compared to real data using the same detection method and measured by Recall and Precision; 4) evaluating spatial crosscorrelation (bivariate Moran’s I), and evaluated using Mantel statistics and cosine similarity to assess how well the simulation reflects the actual cell-type composition.

### Relative performance on data properties and scalability evaluation criteria

An analysis of 13 simulation methods (Fig. 4a) across data properties and scalability evaluations (Fig. 4c & e) revealed variable method performance across metrics. As expected, the performance of the method varied significantly among evaluation metrics, suggesting that there is not a universally effective approach that performs well across data properties and scalability evaluation. We will first examine data property estimation including gene-level, spot-level and spatial level. After that, we will evaluate scalability including measured time and memory in varying numbers of gene and spot, to determine the computational efficiency and feasibility of the simulators for large-scaled datasets.

Interestingly, we observed that certain single-cell simulators such as scDesign2, ZINB-WaVE and SPARsim are equally good as spatial simulators such as scDesign3 and SRTsim in capturing gene-level and spot-level data distributions. Nevertheless, the spatial simulator SRTsim is the only method that consistently excelled in spatial metrics. This is not unexpected, given that spatial simulators are specifically designed to capture spatial relationships. In detail, for gene-level estimation, the performance of methods varied across according to six criteria. Among these, the single-cell methods scDesign2, ZINB-WaVE and SPARsim and the spatial methods SRTsim and scDesign3 (poi) outperformed the others (Fig. 4c). For the other methods, significant differences were noted across the six criteria, with no clear patterns or correlations in their rankings based on each criterion. For spot-level estimation, we also noticed variability in the performance of methods across eight criteria. The single-cell methods scDesign2, ZINB-WaVE and SPARsim and the spatial methods SRTsim and scDesign3 (nb) emerged as top performers (Fig. 4c). It is important to note that SRTsim with the simAdaptor yielded better results than without when assessing scaled variance and mean at the spot-level. For spatial-level estimation, we found that some single-cell simulators, such as SPARsim and Splatter, perform well. They are as effective as certain spatial simulators, such as SRTsim and scDesign3.

Evaluating computational efficiency, each method was tested on subsampled versions of a real dataset (Fig. 4e). Most exhibited good performance, with runtimes under one hour and memory consumption below ten gigabytes (GB). Notably, most single-cell simulators generally excelled in both aspects.

However, a trade-off between efficiency and modeling complexity emerged: ZINB-WaVE achieved top running speeds at the cost of high memory requirements, while scDesign2 demonstrated efficient memory usage but with longer runtimes particularly in scDesign3. The scDesign3 utilizes a copula model to capture correlation, which can be time-consuming, especially as the number of genes scales. In this work, we need to constrain the number of genes (200, 500, 1k) in the dataset to ensure the simulation could be completed within reasonable timeframes. This also highlights the inherent tension between computational demands and the sophistication of the simulation framework. No significant distinction was observed between spatial aware and spatial adaptor approaches in terms of scalability.

### Downstream analytical tasks revealed their relative performance on multi-tasks criteria

Next, we examine spatial tasks such as spatial clustering, spatially variable genes identification, cell type deconvolution, and spatial cross-correlation (Fig. 4d). The objective is to determine the realism of simulated data in subsequent analyses. We observed variability in methods’ performance for four spatial tasks. This also suggests that no single method consistently excels in all spatial task evaluations. In general, most single-cell simulations and spatial simulators perform well in the four spatial tasks evaluated (Fig. 4d). Our evaluation shows that the single-cell methods SPARsim, Splatter, and SymSim as well as the spatial method SRTsim, excel in spatial clustering performance. In the spatial task of cell type deconvolution, we observed that the spatial methods SRTsim and scDesign3 generally outperformed single-cell simulators. For spatially variable gene identification, the single-cell method ZINB-WaVE, along with the spatial methods SRTsim and scDesign3, significantly outperformed others. Additionally, unlike spatial autocorrelation (univariate Moran’s I), which assesses how a single variable correlates with itself across different spatial locations, spatial cross-correlation explores how two distinct variables co-vary spatially. Our analysis demonstrated that the spatial methods SRTsim and scDesign3 (poi) significantly outperformed other simulators in this task.

### Impact of model distribution and dataset characteristic on model performance

Beyond comparing overall method performance, understanding the factors influencing simulation outcomes is crucial for informed method selection and method development. Here we investigate the potential factors influencing simulation results, identifying both the common strengths and weaknesses of current simulation methods, as well as the progress achieved. We first explored the influence of distribution assumptions on model estimates by applying different distributions (Gaussian, Negative binomial, Poisson) to scDesign3. Negative binomial performed the best, followed by Poisson and Gaussian distribution. This observation corroborates the typical distribution modeling approach in the single-cell community, where data are commonly modelled as Negative binomial or Poisson distributions rather than Gaussian. This suggests that capturing overdispersion, a common feature of spatial data, is crucial for accurate modeling. While SPARsim and ZINB-WaVE (utilizing zero-inflated negative binomial and gamma distributions, respectively) maintained good performance, SRTsim excelled across most metrics. This potentially stems from their ability to adaptively select distributions (Poisson, Negative binomial, zero-inflated Poisson and zero-inflated negative binomial) based on the data, offering greater flexibility in capturing complexities.

To examine if the performance of simulation models are consistent across datasets, we examined the KDE test statistics values across various data properties (spot-level, gene-level and spatial-level) on different datasets (Fig. 5). The scDesign3 (nb), SRTsim, and scDesign2 displayed superior consistency across spot-level, gene-level, and spatial-level (Fig. 5a). This is further supported by the forest plot analysis (Fig. 5a) where SRTsim, scDesign3 (nb) and scDesign2 exhibited minimal variability across evaluation metrics. The scDesign3 (gau) was affected by fraction zero, library size and Pearson, potentially due to scDesign3’s Gaussian assumption being unsuitable for sparse data. The Splatter, SymSim, and ZINB-WaVE were significantly influenced by spot-spot Pearson correlation and efficient library size (Fig. 5b). In gene-level, we observed that scDesign3 (gau), Splatter and SymSim’s performance was influenced by higher-order interactions such as mean verse variance and mean verse fraction zero. Most of the single-cell simulators (Splatter, SPARsim, SymSim and ZINB-WaVE) were affected by scaled variance and scaled mean. In spatial-level, we found that most simulators are strongly influenced by how a single variable correlates with itself across different locations. This relationship is measured using Moran’s I. These findings highlight the importance of considering the data type when selecting models and the implementation of a comprehensive collection of data type and evaluation metrics when assessing simulation models.

**Figure 5:**
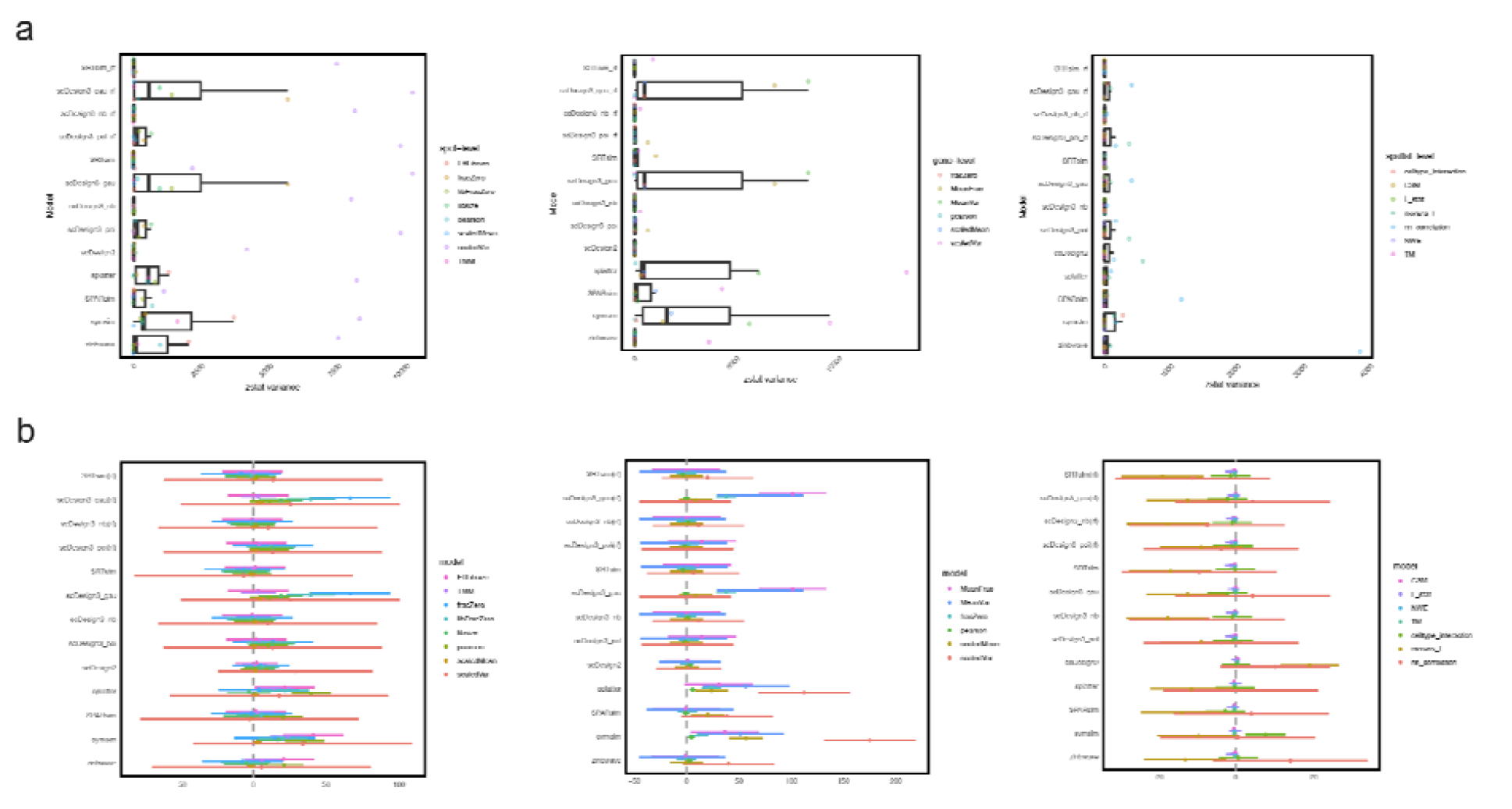
Impact of dataset and evaluation metric on method performance. **a**, Dataset Influence on models, illustrated by a boxplot depicting model-dataset consistency in spot-level, gene-level and spatial-level; **b**, Forest plot representing the impact of submetrics across different models. There are three graphs in each panel representing different evaluation areas, starting from left to right: spot-level, gene-level and spatial-level.

## Discussion

In this study, we presented SpatialSimBench, a multi-task benchmarking study evaluating the performance of overall simulation methods in spatial gene expression data, including eight spatial simulators and five single-cell simulators. Importantly, we introduced a simulation strategy, which we termed simAdaptor approach. We demonstrated that simAdapts enables existing scRNA-seq simulators to simulate spatially resolved data, as well as improving the performance of spatial simulators. We assessed all simulators using ten spatial gene expression datasets with paired single-cell gene expression data and analyzed them across 35 distinct metrics. These metrics covered a range of aspects, including data properties, spatial downstream analytical tasks and scalability. Based on our result, we also explored the effect of distribution assumptions and the consistency of data characteristics on model estimation. Overall, this study provides recommendations for method selection and identifies improvements in future method development.

A major challenge for spatial simulators lies in the heterogeneity of spatial transcriptomics (ST) technologies. For example, image-based ST, such as seqFISH[16], and MERFISH[17], offers high resolution but is limited by a smaller number of target genes. Sequencing-based ST captures expressed RNAs in space but with low spatial resolution, such as ST[18], and 10X Visium[19]. One key finding was the significant impact of sparsity in the spatial transcriptomics data matrix on simulation performance.

Methods like scDesign3 (gau) appear to have a limited effect on higher order interactions. This limitation might be because scDesign3’s underlying assumption of Gaussian distribution may not be appropriate for data that is sparse or has many zeros. Moreover, our analysis further revealed that several single-cell simulators, including SPARsim, Splatter and SymSim, exhibit robust capabilities for handling spatial clustering in downstream analysis. These tools are originally designed for managing complex scenarios inherent and they also retain these features when using simAdaptor.

We also explore the idea that current models for simulating single cells could be slightly modified to analyze spatial gene expression. We raise the question whether a spatially aware simulator adds practical advantage or is it mainly a theoretical justification. In this study, we developed a simAdaptor approach that utilizes existing simulators to generate spatial gene expression data by incorporating spatial segmentation techniques. This approach successfully captured some aspects of real spatial data, including clustering patterns and similar spatial distribution patterns. Yet, when it comes to capturing complex interactions of features across spots and genes, specialized spatial simulators perform better. These spatial simulators prove useful in certain tests, but there is still a lot to gain from integrating single-cell simulation techniques. This raises the question of future method development. Should we modify existing single-cell simulators into spatial versions, or design entirely new structures specifically for spatial data? An interesting possibility lies in combining these approaches, potentially leading to new and powerful methods within the single-cell research community.

Our findings from both SpatialSimBench and simBench studies show remarkable consistency. Both sets of simulations observed that: 1) different methods perform differently depending on the criteria used for evaluation, 2) there is an inherent trade-off between computational efficiency and achieving a good model fit, and 3) the underlying distribution assumptions and datasets used can significantly impact model estimation. While some observations are similar between simBench and SpatialSimBench, there have been significant advancements in the past four years. Notably, simBench studies highlighted that most of the simulation models often underperform higher-order interactions metrics. Conversely, SpatialSimBench studies demonstrate substantial improvement in both spatial and single-cell simulators. More importantly, SpatialSimBench establishes a novel benchmarking framework that bridges the gap in single-cell RNA sequencing simulations by incorporating spatial context. This innovation emphasizes the potential for creative benchmarking approaches in the single-cell community, even with ongoing method development, as these methods can still be significantly influenced by the chosen evaluation framework. In essence, both studies confirm the importance of considering evaluation criteria, computational cost, and underlying assumptions when selecting methods. However, SpatialSimBench takes a significant leap forward by introducing a creative benchmarking strategy that integrates spatial context, paving the way for future advancements in the field.

Recently, the Open Problems in Single-Cell Analysis [20] offers an open-source, community-driven platform for benchmarking various formalized tasks in single-cell analysis. It covers a broad range of areas, including batch integration, cell-cell communication inference, and spatial decomposition contributed by community effort. In developing interactive software for SpatialSimBench, we aim to contribute to the broader single-cell analysis community through the Open Problems Framework and also to offer a more detailed and interactive experience through developing our own Shiny app.

In summary, our work offers valuable insights for both benchmark and method development in spatial transcriptomics. We demonstrated the usefulness of our framework by summarizing how different methods perform across various aspects. This can help users choose the appropriate method for their needs and also pinpoint areas where developers can improve existing methods. Furthermore, we’ve introduced a novel benchmark evaluation framework in spatial simulation by integrating spatial information into traditional single-cell RNA sequencing simulations. Our findings are a valuable resource for both biologists seeking to analyze their spatial transcriptomics data and method developers aiming to advance the state-of-the-art.

## Material & Methods

### Data description

In this benchmark study, we collected a total of ten spatial transcriptomics datasets alongside reference scRNA-seq datasets. The spatial transcriptomics datasets include a range of protocols, tissue types, and health conditions, from human and mouse. The scRNA-seq datasets were obtained by 10x Chromium and Smart-seq platforms with all cell type labels being accessible publicly. Details of all datasets are available in Supplementary Table 1.

Regarding data preprocessing, we normalized the expression matrix for spatial transcriptomics datasets and scRNA-seq datasets by scater[21]. In consideration of time constraints and the convention in single-cell research that usually considers a selection of 1,000 to 2,000 genes adequate for downstream analysis, we selected the top 1,000 spatially variable genes when the total number of genes exceeded 1,000; otherwise, all genes were included. For the scRNA-seq datasets, our focus was on ensuring data integrity, by following a stringent quality control process[22]. Following the QC metric from Seurat [23], we filtered cells with unique feature counts over 2,500 or less than 200 and larger than 5% mitochondrial counts.

To estimate cell-type composition, we employed BayersSpace as it outperforms on cell type clustering methods for spatial transcriptomics data[24]. CARD integrated cell-type-specific information from scRNA-seq data with correlation in cell-type composition across tissue location, achieved through the implementation of a Conditional Autoregressive (CAR) modeling approach.

### Benchmark method

Our study introduces a novel approach that integrates spatial context into traditional single-cell RNA sequencing (scRNA-seq) simulation models. We termed this approach simAdaptor. This involves a two-step process. Initially, spatial data are clustered to identify regions with similar expression profiles. This is done using BayersSpace, a method designed specifically for spatial clustering. After clustering, we applied established simulation methods to simulate the data separately in each cluster. This process enables us to indirectly incorporate spatial variables into existing simulation models. In comparison, the non-simAdaptor approach applies spatial simulation onto each dataset directly without modification.

Our literature review identified six spatial simulators: scDesign3[25], SRTsim[26], scMultiSim[9], stLearn[3], Spider[7]and spaSim[4]. While scMultiSim employs cell differentiation trees and gene regulatory networks to simulate spatial gene expression data at the cell level, this approach does not take gene expression as the input. It is thus not possible to assess whether this method performs well on gene-level, spot-level and spatial-level data properties. On the other hand, Spider and stLearn process scRNA-seq data to simulate count data along with spatial locations. However, their generated spatial data are challenging to evaluate. Specifically, the absence of ground truth spatial location information in their outputs makes it difficult to compare with spatial patterns. SpaSim only stimulates cell locations by simulated image without spatial data. Therefore, we excluded these four spatial simulators for two primary reasons: the unsuitability of their input formats in capturing both spatial and gene expression details, and the inherent difficulties in comparing simulated spatial patterns without verifiable spatial location information. To ensure a focus on spatial data generation with a fair comparison, we selected scDesign3 and SRTsim. To compare how spatial simulators compare with existing single-cell simulators, We further expanded our analysis to include well-performing single-cell simulators from the SimBench study, such as Splat[12], ZINB-WaVE[11], SymSim[13], scDesign2[27], and SPARsim[10]. All of the simulation models are implemented in R. Comprehensive information about these methods, including code versions, associated publications, and default parameter configurations, can be found in Supplementary Table 2.

### Evaluation of data properties

We developed a unified pipeline to assess the efficacy of the simulation models. Within this pipeline, the first evaluation is data properties including spot-level, gene-level and spatial level. After computing the metric for both the real and simulated datasets, we employ density plots and KDS scores for each metric to assess the similarity between the real and simulated datasets.

#### Spot-level

- Library size: total sum of UMI counts across all genes
- TMM: weight trimmed mean of M-values normalization factor[28]

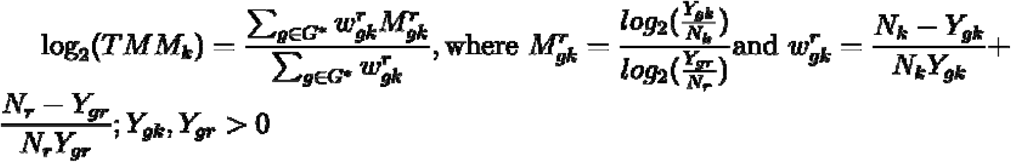

where ***Y***_***gk***_ is the observed count for gene ***g*** in library ***k*** summarized from the raw reads and ***N***_***k***_ is the total number of reads for library ***k***; ***Y***_***gk***_ is the observed count for gene ***g*** in library ***τ*** summarized from the raw reads; ***N***_***r***_ is the total number of reads for library ***τ***
- Effective library size: library size multiplied by TMM
- Scaled variance: z-score standardization of the variance of expression matrix in terms of log2 CPM

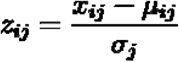

where ***x***_***i***_***j*** is the ***log***_**2**_***CPM*** value for the ***i***^***th***^ gene in the ***j***^***th***^ sample; ***μ***_***j***_ is the mean of the ***log***_**2**_***CPM*** values for the ***j***^***th***^ sample across all genes; ***σ***_***j***_ is the standard deviation of the ***log***_**2**_***CPM*** values for the ***j***^***th***^ sample across all genes
- Scaled mean: z-score standardization of the mean of expression matrix in terms of log2 CPM

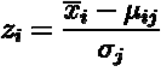

where 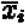 is the mean of the ***log***_**2**_***CPM*** value for the ***i***^***th***^ gene across all samples; ***μ*** is the mean of the overall average ***log***_**2**_***CPM*** values for samples across all genes; ***σ*** is the standard deviation of mean across all genes
- Fraction zero: fraction zero per spot
- Library size vs fraction zero: the relationship between library size and the proportion of zero per gene
- Sample Pearson correlation

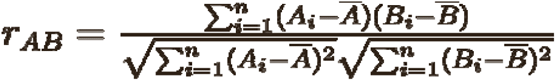

where ***A***_***i***_ and ***B***_***i***_ are the ***log***_**2**_***CPM*** values of gene ***A*** and ***B*** in the ***i***^***th***^ sample; ***Ā***_***i***_ and 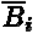 are the mean ***log***_**2**_***CPM*** values of gene ***A*** and ***B*** across all samples

#### Gene-level

- Fraction zero gene: proportion of zero per gene
- Scaled variance: z-score standardization of the variance of expression matrix in terms of log2 CPM

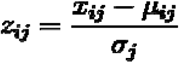

where ***x***_***i***_***j*** is the ***log***_**2**_***CPM*** value for the ***i***^***th***^ sample in the ***j***^***th***^ gene; ***μ***_***j***_ is the mean of the ***log***_**2**_***CPM*** values for the ***j***^***th***^ gene across all samples; ***σ***_***j***_ is the standard deviation of the ***log***_**2**_***CPM*** values for the ***j***^***th***^ gene across all samples
- Scale mean: z-score standardization of the mean of expression matrix in terms of log2 CPM

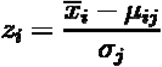

where 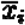 is the mean of the ***log***_**2**_***CPM*** value for the ***j***^***th***^ samples across all genes; ***μ*** is the mean of the overall average ***log***_**2**_***CPM*** values for genes across all samples; ***σ*** is the standard deviation of mean across all samples.
- Mean vs variance: the relationship between mean expression and variance expression
- Mean vs variance (scale): the scale of the relationship between mean expression and variance expression by log-normal transformation
- Mean vs fraction zero: the relationship between mean expression and the proportion of zero per gene
- Gene Pearson correlation

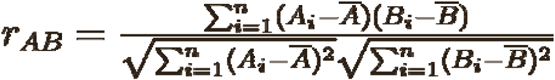

where ***A***_***i***_ and ***B***_***i***_ are the ***log***_**2**_***CPM*** values of sample ***A*** and ***B*** in the gene; ***Ā***_***i***_ and 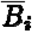 are the mean ***log***_**2**_***CPM*** values of sample ***A*** and ***B*** across all genes.

#### Spatial-level

This section of our study focuses on spatial-level metrics. We used transition matrix, neighborhood enrichment matrix and centralized score matrix to depict the global spatial patterns evident in spatial transcriptomic data, originating from spider[7]. The other four metrics (cell type interaction, Moran’s I, L statistic and nearest neighbor correlation), originating from scFeatures[14], are specifically designed for a multi-view representation of spatial data and includes feature types commonly used in spatial analysis.

- Transition Matrix (TM): This is a normalized matrix representing transition frequencies, which embodies the probabilities of transitioning from one state to another within a Markov chain framework. In this context, the TM elucidates the interrelationships among spatial clusters in each space. Each element in the matrix signifies the transition probability from one spatial cluster to another, thereby mapping the dynamic interplay of spatial clusters
- Neighborhood enrichment matrix (NEM): This matrix quantifies the enrichment observed between each pair of spatial clusters. It serves to systematically assess the interaction between different clusters within a spatial context, providing insights into the relative connectivity between various spatial clusters

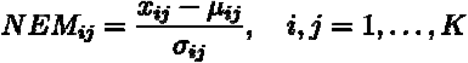

where ***x***_***i***_***j*** is the number of connection between ***C***_***i***_ and ***C***_***j***_; ***μ***_***ij***_ is the expected mean; ***σ***_***ij***_ is the standard deviation
- Centralized score matrix (CSM)

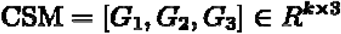

where ***G***_**1**_ is group degree centrality; ***G***_**2**_ is the average clustering coefficient; ***G***_**3**_ is the group closeness centrality. Each of the details show below:
  ∘ Group Degree Centrality: calculates the ratio of spots within one spatial cluster that are connected to spots in another spatial cluster. It assesses the inter-cluster connectivity, indicating the extent to which one cluster is interlinked with another

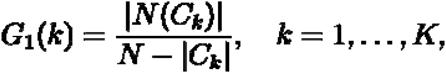

where ***C***_***k***_ is the group of ***k***^***th***^spatial cluster; ***N*(*C***_***k***_)is the neighbors of all the spot in ***C***_***k***_; ***N*** is the number of spots
  ∘ Average Clustering Coefficient: measures the propensity for a spot within a spatial cluster to be connected to spots in another cluster. It provides insights into the likelihood of inter-cluster associations, reflecting the tendency of spots to form connections beyond their immediate cluster

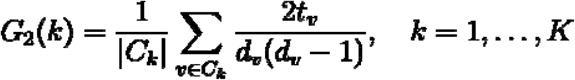

where ***t***_***v***_ is the number of triangles around spot ***v***; ***d***_***v***_ is the degree of spot ***v***, which is the number of connections or edges that spot ***v*** has
  ∘ Group Closeness Centrality: the normalized inverse sum of distances from a spatial cluster to all spots in a different spatial cluster. It quantifies the relative proximity or accessibility of one cluster to all spots in another, offering a measure of how closely or centrally positioned a cluster is concerning another cluster in the spatial arrangement

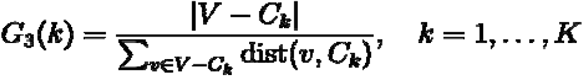

where ***V*** is the number of spot; **dist(*v***,***C***_***k***_)is the shortest distance between spatial cluster ***C***_***k***_ and spot ***v***.
- Cell type interaction: assume the nearest neighbors should be the cells captured within each spot and consider them as the spatial interaction pairs. Then used the estimated cell type proportion in each spot to calculate the spatial interaction between cell types. For details refer to scFeatures.
- Moran’s I: measure spatial autocorrelation, meaning how strongly the feature expression value in a sample cluster or disperse.

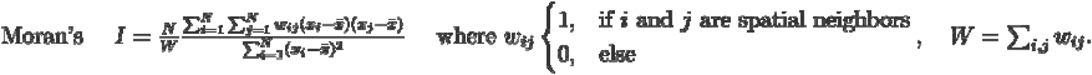

where ***x***_***i***_ and ***x***_***j***_ are the gene expression values of ***i***^***th***^ cell and ***j***^***th***^ cell; 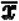 is the average gene expression value of one gene; ***N*** is the number of cells
- L statistics: the L value between the pairs of genes by estimation cell type proportion. For details refer to scFeatures
- Nearest neighbor correlation: Pearson correlation for gene expression between a spot with its nearest neighbor spot. For details refer to scFeatures.

### Evaluation of spatial downstream analysis

This evaluation is spatial downstream analysis including spatial clustering, cell type deconvolution, spatially variable gene identification and spatial cross-correlation. We performed each algorithm on the real experimental dataset and simulated dataset and compared the similarity of the result.

#### Spatial Clustering

Spatial clustering refers to cluster or group spots based on similar expression patterns across a spatial domain. Here we use the `BayersSpace` package to perform spatial clustering. We apply Adjusted rand index (ARI) and Normalized mutual information (NMI) to evaluate the spatial clustering result between real data and simulated data.

- Adjusted rand index (ARI): measure the similarity between two clusters in real and simulated datasets

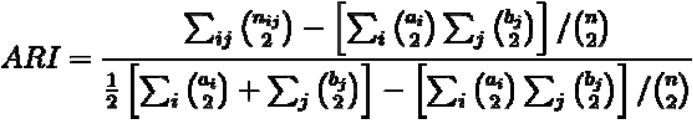

where ***n*** is the total number of spots; ***a***_***i***_ is the number of items in the ***i***^***th***^ spatial cluster of the spatial clustering in real dataset; ***b***_***j***_ is the number of items in the ***j***^***th***^ spatial cluster of the spatial clustering in simulated dataset; ***n***_***ij***_ is the number of items that are in both ***j***^***th***^ the cluster of the spatial clustering in real dataset and the ***j***^***th***^ cluster of the spatial clustering in a simulated dataset.
- Normalized mutual information (NMI): a measure of the mutual dependence between the real and simulated spatial clusters.

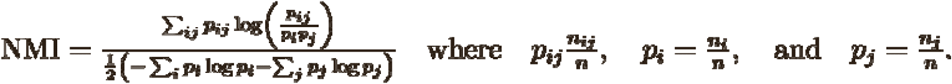

where ***n*** is the total number of spots; ***n***_***i***_ is the number of items in the ***i***^***th***^ spatial cluster of the spatial clustering in real dataset; ***n***_***j***_ is the number of items in the ***j***^***th***^ spatial cluster of the spatial clustering in simulated dataset; ***n***_***ij***_ is the number of items that are in both the ***i***^***th***^ cluster of the spatial clustering in real dataset and the ***j***^***th***^ cluster of the spatial clustering in a simulated dataset.

#### Cell type deconvolution

Cell type deconvolution refers to interpreting mixed signals within tissue compartments to identify proportions of cell types per spot. This is only relevant to spot-based technology where you have multiple cells per spot. However, this is not relevant for a single-cell based platform where you are able to measure the expression for a single cell. Here we use `CARD` package to perform cell type deconvolution.

We assume that the number of genes per spot is ***J*** and the number of spot is ***I*** in spatial transcriptomics data; ***X***_***ij***_ represent the spatial gene expression value of gene ***j*** in the ***i*** spot; ***T***_***ik***_ and ***P***_***ik***_ are the true and predicted proportion of cell type ***k***.

- RMSE: root mean square deviation is calculated between ***T***_***ik***_ and ***P***_***ik***_ of per cell type. After that, we normalize them by sum of proportions among all the spots ***S***_***k***_. Then, we average all the RMSE as final RMSE

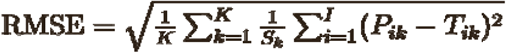
- JSD: Jensen-Shannon divergence was calculated between ***T***_***k***_ and ***P***_***k***_ per cell type in all spots. We use Kullback–Leibler divergence (KL) to calculate JSD. ***Q*(*T***_***k***_) and ***Q*(*P***_***k***_) is true distribution and algorithm-predicted of cell type ***k***. Then, we average all the JSD as final JSD

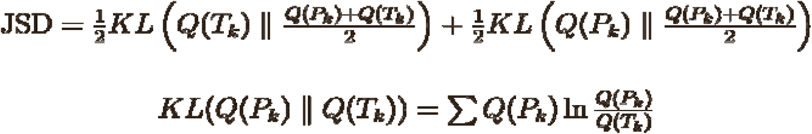

#### Spatially variable genes (SVG) identification

Spatially variable genes identification refers to identifying genes whose expression levels vary significantly across different spatial coordinates or regions. Here we use SPARK-X package to perform SVG identification. We apply Precision and Recall to evaluate the results.

- Precision: the proportion of correctly identified items in simulated datasets

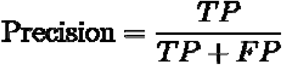

where false positive (FP) is the number of genes that are incorrectly identified as SVG when they are not SVG; true positive (TP) is the number of genes that are correctly identified as SVG
- Recall: the proportion of real SVG correctly identified in the simulated dataset

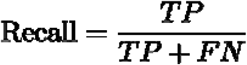

where false negative (FN) is the number of genes that are incorrectly predicted as non-SVG when they are true SVG

#### Spatial cross-correlation

Spatial cross-correlation explores how two distinct genes co-vary spatially by bivariate Moran’s I.

- Bivariate Moran’s I: Moran’s I between genes and was calculated by the following.

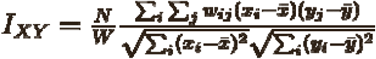

where ***w***_***ij***_ represents a spatial weight matrix and; ***w***_***ij***_ **= 0**,***W* = ∑ *w***_***ij***_**; *N*** represents the number of spots.
- Cosine similarity: measure similarity between bivariate Moran’s I of real dataset ***A*** and that of in simulation dataset ***B***.

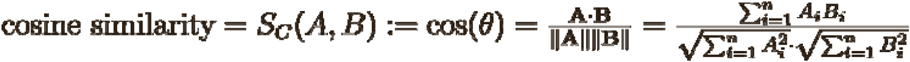

where ***A***_***i***_ and ***B***_***i***_ are the ***i***^***th***^ components of vectors ***A*** and ***B***, respectively.
- Mental statistics: The test statistic for the Mantel test, which is a correlation coefficient calculated between bivariate Moran’s I of real dataset ***A*** and that of in simulation dataset ***B***

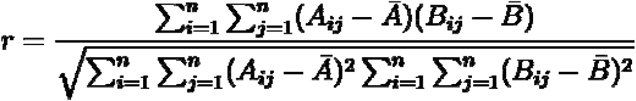

where ***A***_***ij***_ is the element in the ***i***^***th***^ spot and ***j***^***th***^ gene of ***A***; ***B***_***ij***_ is the element in the ***i***^***th***^ spot and ***j***^***th***^ gene of ***B***; ***n*** is the number of spots in ***A*** or ***B***

### Evaluation of method comparison in each score and rank-based overall score

In order to summarize the results derived from multiple datasets and criteria, we employed a multi-step approach to generate scores. This process was essential due to the utilization of distinct metrics, with KDE scores being applied for both spot-wise and gene-wise evaluations, as well as a part of the spatial pattern assessments. Also, spatial clustering and some spatial pattern evaluation utilized different scoring systems, necessitating a structured method to integrate these diverse scores coherently. Details show below:

In a kernel density estimation (KDE) test, the null hypothesis assumes that the two estimated densities are identical. The integrated squared error (ISE) quantifies the discrepancy between these estimates. Under the null hypothesis, the final test statistic is calculated based on the ISE:

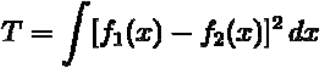

where ***f***_**1**_ and ***f***_**2**_ are the kernel density estimates of sample 1 and sample 2, respectively. The implementation from the R package ks (v1.14.2) was used for the KDE test performed in this study.

To quantify the similarity between the simulated and real distributions, we leveraged the test statistic from the KDE test. We transformed this statistic into a similarity measure ranging from 0 (completely dissimilar) to 1 (perfectly similar) using the below equation. This transformation involved scaling the absolute value of the statistic to lie between 0 and 1, and then inverting it (subtracting from 1).

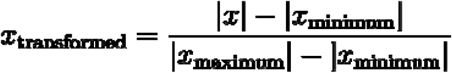

where ***x*** is the raw value before transformation. This transformation is applied to the KDE scores obtained from all methods across all datasets. To ensure the transformation works consistently, ***x***_***minimum***_ and ***x***_***maximum***_ are defined based on the range of all these KDE scores. The goal of this transformation is to follow the principle of “higher is better,” making the resulting values easier to interpret.

Our initial step involved generating individual scores for each dataset via each simulation model. This allowed us to summarize the performance of each method across all datasets by calculating their average scores. This resulted in a single score of comparison among all methods.

Following this, we established an overarching score for each method by integrating the metrics to each method’s evaluation across datasets. For instance, in calculating a rank-based overall score focused on spot-wise evaluation, we first computed the KDE scores for each method across nine metrics. Then, we arranged the spot-wise metrics of these methods in ascending order—effectively ranking them from best to worst—across nine metrics. The final step involved averaging these rankings to obtain an overall accuracy score, as follow:

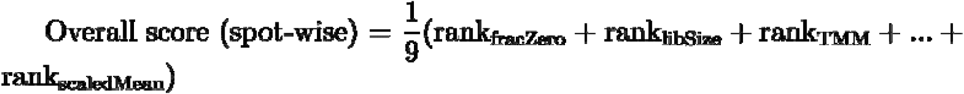

### Evaluation of impact of dataset distribution in method performance

To synthesize the outcomes from various criteria across four assessments, we adopted a layered method to calculate scores for every dataset. Initially, we divided the process into four distinct evaluations: spot-wise, gene-wise, spatial clustering, and spatial patterns, recognizing that each assessment captures unique aspects.

In the subsequent phase, we computed the average of scores from several metrics for each dataset within every evaluation. We then determined the overall score for each dataset across the simulation model in all four evaluations. For instance, in assessing the consistency of a dataset within the spot-wise evaluation, we averaged the KDE score from nine different criteria for a single dataset using one method.

### Evaluation of impact of model distribution in method performance

To assess the impact of models within a specific metric, we performed linear regression analyses for each metric separately. This approach allowed us to quantify the effect of switching from one model to another within the same metric context. By comparing the regression coefficients, we could determine which models had statistically significant impacts on the scores, thereby evaluating each model’s effectiveness across different metrics. This analysis was conducted using the lm function in the built-in stats package in R.

For each metric, a linear regression model was fitted where the dependent variable was a KDE score, and the independent variable was the type of model with the formula defined as:

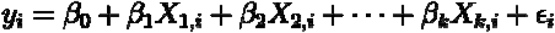

where ***y***_***i***_ represents score for the ***i***^***th***^ observation; ***X***_***1***_,_***i***_,***X***_***2***_,_***i***_,***X***_***k***_,_***i***_ are indicator variables (dummy variables) derived from the model factor, representing the presence (1) or absence (0) of each category (excluding the reference category) for the ***i***^***th***^ observation; ***β***_**0**_ is the intercept term and ***β***_**1**_, ***β***_**2**_,**… *β***_***k***_are the regression coefficients for each of the indicator variables.

### Evaluation of scalability

To mitigate potential confounding effects, our analysis of scalability was confined to Dataset 7, which we systematically downsampled to produce datasets with varying numbers of spots and genes, specifically including spot counts of 200, 500, 1,000, 3,000, and 5,000, and gene counts of 200, 500, and 1,000, resulting in 15 downsampled datasets.

The execution time for each method was gauged using the Sys.time function in R and the time.time function in Python. Tasks failing to complete within the allotted time frame were deemed to have generated no results. To capture the peak memory usage of R methods, we utilized the psutil package to monitor the maximal Resident Set Size, with all measurements taken thrice and averaged for accuracy.

In the simAdaptor approach, we implemented a two-step simulation process: initially conducting spatial clustering on the dataset and labeling each spot, followed by sequentially integrating each spatial cluster into the simulation models. The duration and memory consumption for both stages were recorded separately and are presented in Supplementary Figure 2. Conversely, for one-step approaches where the entire dataset is fed into the simulation model simultaneously, we only tracked the time and memory requirements of this singular process.

Computational resources for these tests included running the 13 simulation methods on an R server equipped with Intel Core i9-14900K CPUs (5.2 GHz, 36 MB Smart Cache, and a total of 24 CPU cores) and 64 GB of DDR5 6000 MHz memory.

## BACKMATTER

### Ethics statement

The authors declare no competing interests.

## Data Availability

All datasets used in this study are publicly available. Details on each datasets, including their accession ID are provided in Supplementary Table 1.

### Acknowledgments

The authors thank all their colleagues, particularly at the Sydney Precision Data Science Centre, Charles Perkins Centre for their support and intellectual engagement.

## Funding

This work is supported by the AIR@innoHK programme of the Innovation and Technology Commission of Hong Kong, Judith and David Coffey funding and Chan Zuckerberg Initiative Single Cell Biology Data Insights grant (DI2-0000000197) to YC and JYHY; NHMRC Investigator APP2017023 to JYHY. The funding source had no role in the study design, in the collection, analysis, and interpretation of data, in the writing of the manuscript, or in the decision to submit the manuscript for publication.

## Author Contribution

JYHY and YC conceived and led the study with design input from XL. XL ran the benchmarking studies and performed interpretation of the evaluation framework with close guidance from JYHY and YC. All authors contributed to the writing, editing, and approval of the manuscript.

## Notes

### Competing Interest Statement

The authors have declared no competing interest.

