## Supplement for "Multi-task benchmarking of spatially resolved gene expression simulation models"

### Supplementary Figures

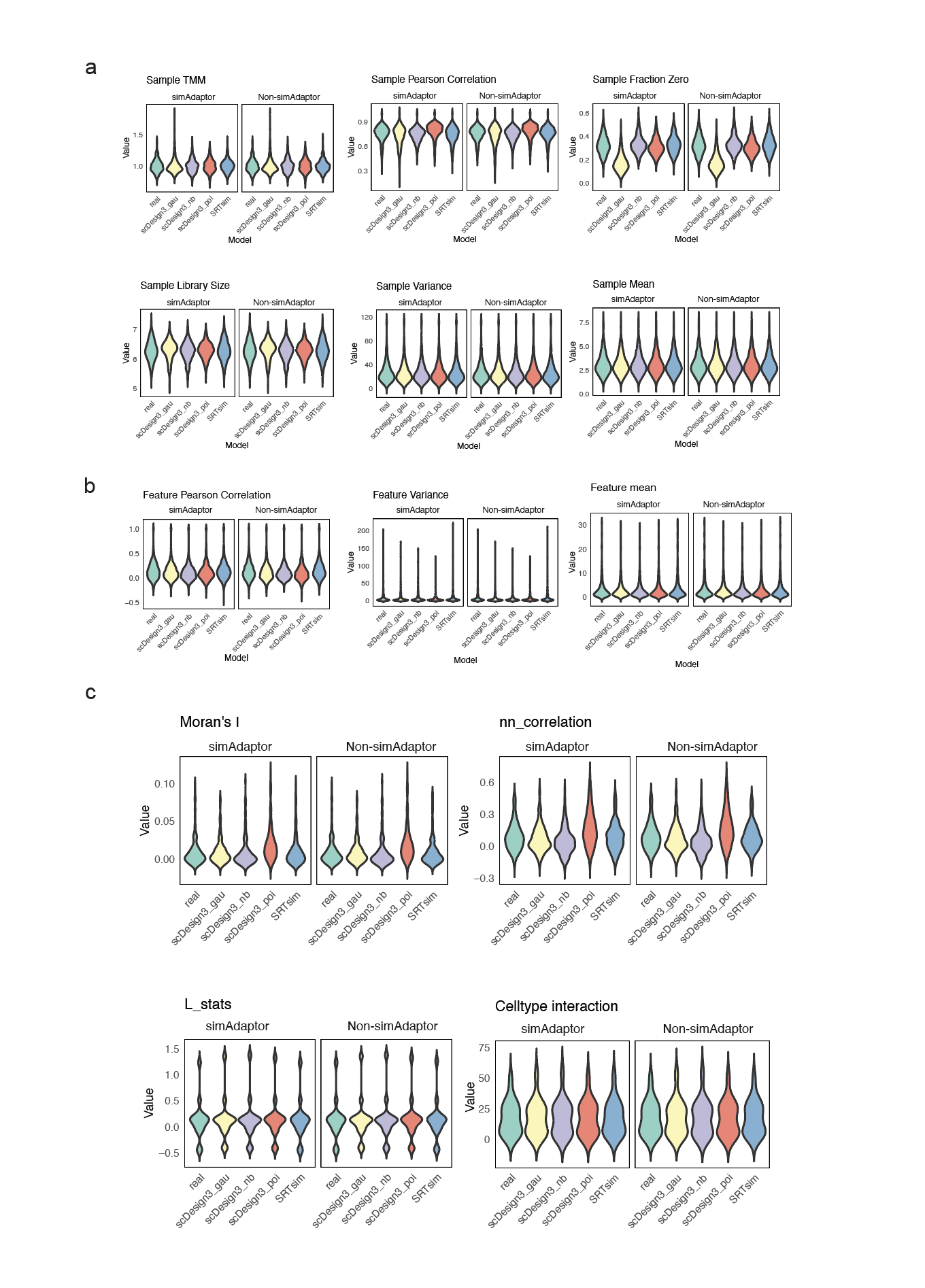

**Supplementary Figure 1. Comparison of approaches based on using simAdaptor and those without simAdaptor, with an assessment of data properties results from spatial simulators.** **a**, Visualize the real and simulation in boxplot across spot-level metrics. **b**, Visualize the real and simulation in boxplot across gene-level metrics. **c**, Visualize the real and simulation in boxplot across spatial-level metrics.

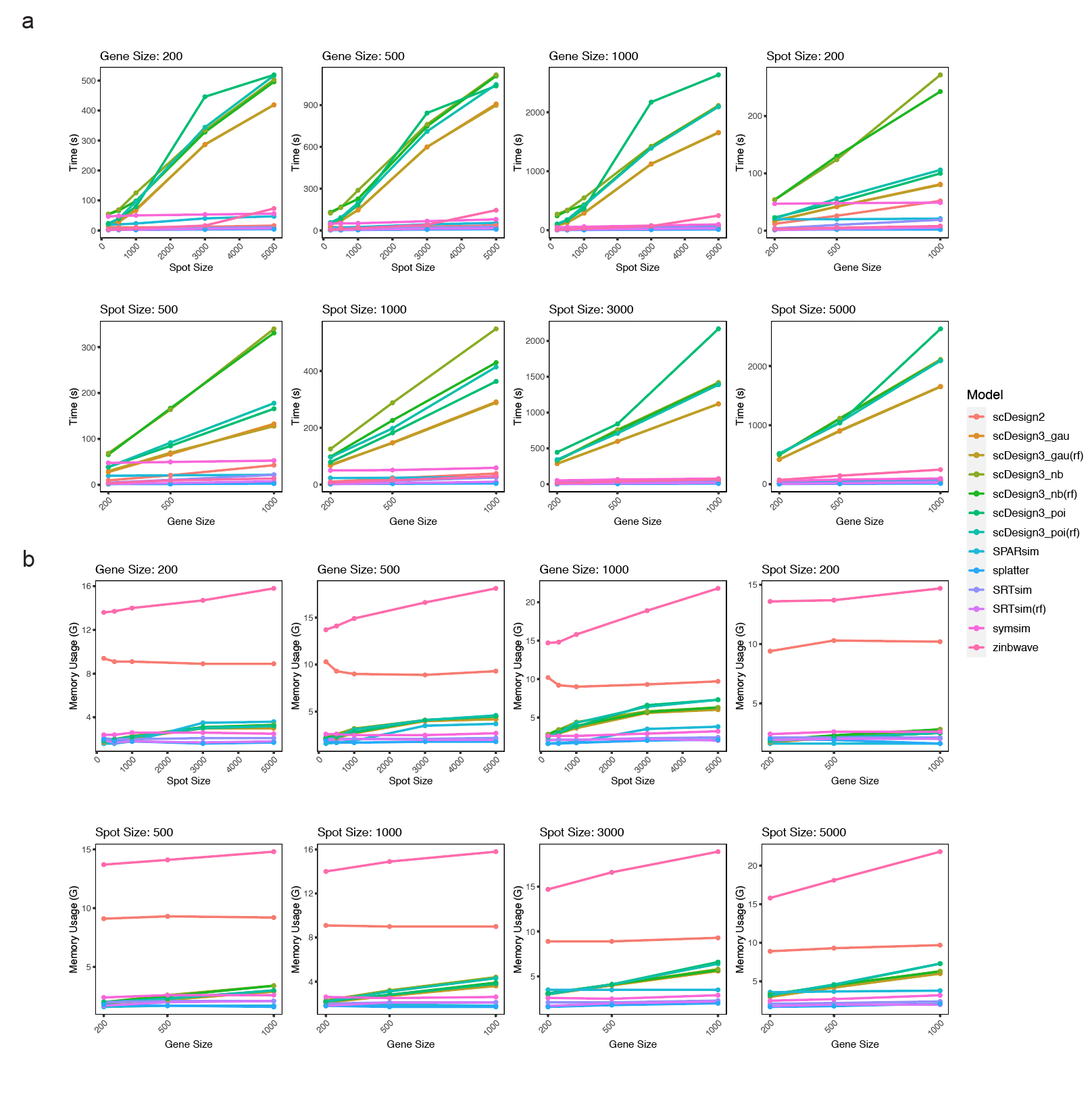

**Supplementary Figure 2: Run time and memory consumption of each method across various spot sizes and gene sizes.** **a**, Runtime of each method across spot size and gene size. **b**, Maximal memory usage of each method across spot size and gene size.

### Supplementary Tables

**Supplementary Table 1.** Details of the spatial gene expression data and single-cell data evaluated in this study. The first table is spatial gene expression data and the second table is single-cell data.

| Dataset | Data modality | species | tissue | health state | protocol | Spot / cell number | Gene number | Source | Data source paper |
| --- | --- | --- | --- | --- | --- | --- | --- | --- | --- |
| Dataset 1 | Spatial | Human | breast | cancer | Visium | 4744 | 28402 | CID3586 [GSE176078](https://www.ncbi.nlm.nih.gov/geo/query/acc.cgi?acc=GSE176078) | A single-cell and spatially resolved atlas of human breast cancers [[29]](https://paperpile.com/c/6v96r1/mqTr) |
|  | Single cell | Human | breast | cancer | Chromium | 6178 | 21164 |  |  |
| Dataset 2 | Spatial | Human | osteosarcoma | normal | MERFISH | 645 | 12903 | [Link](https://www.pnas.org/doi/suppl/10.1073/pnas.1912459116/suppl_file/pnas.1912459116.sd12.csv) | Spatial transcriptome profiling by MERFISH reveals subcellular RNA compartmentalization and cell cycle-dependent gene expression. [[30]](https://paperpile.com/c/6v96r1/MiAu) |
|  | Single cell | Human | osteosarcoma | normal | Chromium | 9234 | 19098 |  |  |
| Dataset 3 | Spatial | Human | prostate | normal | Visium | 277 | 36601 | [GSE159697](https://www.ncbi.nlm.nih.gov/geo/query/acc.cgi?acc=GSE159697) | Vitamin D sufficiency enhances differentiation of patient-derived prostate epithelial organoids [[31]](https://paperpile.com/c/6v96r1/XmqA) |
|  | Single cell | Human | prostate | normal | Chromium | 4740 | 27400 |  |  |
| Dataset 4 | Spatial | Mouse | brain | normal | Visium | 2577 | 31053 | [Link](https://github.com/BayraktarLab/cell2location) | Cell2location maps fine-grained cell types in spatial transcriptomics [[32]](https://paperpile.com/c/6v96r1/kZGe) |
|  | Single cell | Mouse | brain | normal | Chromium | 40532 | 12820 |  |  |
| Dataset 5 | Spatial | Mouse | Fibrosarcoma | tumor | Visium | 2125 | 15976 | Link | Multi-resolution deconvolution of spatial transcriptomics data reveals continuous patterns of inflammation [[33]](https://paperpile.com/c/6v96r1/CVVs) |
|  | Single cell | Mouse | Fibrosarcoma | tumor | Chromium | 7185 | 25069 |  |  |
| Dataset 6 | Spatial | Human | breast | cancer | Visium | 4744 | 28402 | [Github](https://github.com/CaiGroup/seqFISH-PLUS) | Transcriptome-scale super-resolved imaging in tissues by RNA seqFISH [[16]](https://paperpile.com/c/6v96r1/JWU3) |
|  | Single cell | Human | breast | cancer | Smart-seq | 14249 | 34041 |  |  |
| Dataset 7 | Spatial | Mouse | gastrulation | normal | seqFISH | 8425 | 351 | [Link](https://content.cruk.cam.ac.uk/jmlab/SpatialMouseAtlas2020/) | Integration of spatial and single-cell transcriptomic data elucidates mouse organogenesis [[34]](https://paperpile.com/c/6v96r1/YvnA) |
|  | Single cell | Mouse | gastrulation | normal | Chromium | 4651 | 19103 |  |  |
| Dataset 8 | Spatial | Mouse | olfactory bulb | normal | ST | 278 | 182 | [Link](https://www.science.org/doi/10.1126/science.aaf2403) | Visualization and analysis of gene expression in tissue sections by spatial transcriptomics [[18]](https://paperpile.com/c/6v96r1/pYOc) |
|  | Single cell | Mouse | olfactory bulb | normal | Chromium | 12640 | 182 |  |  |
| Dataset 9 | Spatial | Mouse | hindlimb muscle | normal | Visium | 884 | 982 | [GSE161318](https://www.ncbi.nlm.nih.gov/geo/query/acc.cgi?acc=GSE161318) | Strength in numbers: Large-scale integration of single-cell transcriptomic data reveals rare, transient muscle progenitor cell states in muscle regeneration [[35]](https://paperpile.com/c/6v96r1/JaSb) |
|  | Single cell | Mouse | hindlimb muscle | normal | Chromium | 4748 | 14360 |  |  |
| Dataset 10 | Spatial | Human | pancreatic ductal adenocarcinomas | normal | ST | 428 | 25753 | [GSE111672](https://www.ncbi.nlm.nih.gov/geo/query/acc.cgi?acc=GSE111672) | Integrating microarray-based spatial transcriptomics and single-cell RNA-seq reveals tissue architecture in pancreatic ductal adenocarcinomas [[36]](https://paperpile.com/c/6v96r1/MhWw) |
|  | Single cell | Human | breast | cancer | Chromium | 1926 | 19736 |  |  |

**Supplementary Table 2.** Details of the spatial and single-cell simulation methods evaluated in this study.

| Methods | Year of publication | package version | parameter setting | spatial information required |
| --- | --- | --- | --- | --- |
| scDesign3_nb | 2023 | 1.1.1 | family_use = 'nb', others are default | Yes |
| scDesign3_poi | 2023 | 1.1.1 | family_use = 'poi', others are default | Yes |
| scDesign3_gau | 2023 | 1.1.1 | family_use = 'gaussian', others are default | Yes |
| SRTsim | 2023 | 0.99.6 | default | Yes |
| Splatter | 2017 | 1.26.0 | default | No |
| ZINB-WaVE | 2018 | 1.24.0 | default | No |
| SymSim | 2019 | 0.0.0.9000 | default | No |
| scDesign2 | 2021 | 0.1.0 | default | No |
| SPARsim | 2020 | 0.9.5 | default | No |
